## SupplementaryData for "Arthropod IGF, Relaxin and Gonadulin, putative orthologs of *Drosophila* insulin-like peptides 6, 7 and 8, likely originated from an ancient gene triplication"

by Jan A. Veenstra

The supplementary data for this manuscript consists of three elements: this pdf document and two spreadsheets.

Spreadsheet 1 contains contains protein sequences for various insulin-like peptides and their putative precursors, as well as their genbank accession numbers or the terms Artemis or Trinity. Artemis indicates that the sequence in question was derived from a genome assembly, Trinity that the sequence was obtained using the Trinity program on a collection of transcriptome SRAs. In a few instances a combination of Trinity and Artemis was used to obtain a sequence.

Spreadsheet 2 contains the expression of some of these genes for a more limited number of species (there is little sense if the only transcriptome SRAs available for a species are from whole animals of unknown sex and unknown physiological status, or when then there are only very few such SRAs). The data contain the SRA identifier, the number of spots of each SRA and then for each protein of interest two numbers, one in blue, which is number of half spots that contains coding sequence of the protein of interest. The second number in black is the blue number multiplied by 1,000,000 and divided by the number of total spots, which yields a relative number.

The content of the remainder of this document is listed below:

The SRAs used in this study page 2

Suppl. Fig. 1. The detailed sequence similarity tree of arthropod ilps. page 6

Suppl. Fig. 2. Decapod gonadulin 2 precursors with four disulfide bridges. page 10

Suppl. Fig. 3. Gene structures of aIGF genes from *Aedes, Sitobion* and *Oncopeltus*. page 11

Suppl. Fig. 4. Altenative splicing of *Apis mellifera* aIGF. page 12

Suppl. Fig. 5. Details of altenative splicing of *Bombyx mori* aIGF. page 13

Suppl. Fig. 6. Sequence alignment of some dipteran aIGFs. page 14

Suppl. Fig. 7. Sequence alignment of decapod relaxins. page 15

### SRAs used:

***Apis mellifera*:**
SRR035881, SRR035924, SRR1182528, SRR1182529, SRR1182578, SRR1182579, SRR1182580, SRR1182581, SRR1182637, SRR1182638, SRR1182639, SRR1182640, SRR1182641, SRR1182642, SRR1239302, SRR1239303, SRR1239304, SRR1239305, SRR1239306, SRR1239307, SRR1239308, SRR1239309, SRR1239310, SRR1239311, SRR1239312, SRR1239313, SRR1254946, SRR1254947, SRR1254948, SRR1254950, SRR1254951, SRR1254952, SRR1254954, SRR1254956, SRR1254957, SRR1254958, SRR1254959, SRR1254960, SRR1255009, SRR1255010, SRR1255011, SRR1255012, SRR1255013, SRR1255014, SRR1255064, SRR1255065, SRR1255066, SRR1255068, SRR1255149, SRR1255150, SRR1255151, SRR1255152, SRR1255153, SRR1255154, SRR1255260, SRR1255326, SRR1255456, SRR1255541, SRR1255542, SRR1255543, SRR1255544, SRR1255545, SRR1255546, SRR1269199, SRR1613229, SRR2153227, SRR2153228, SRR2153229, SRR2153257, SRR2153259, SRR2153260, SRR2153280, SRR2153281, SRR2153282, SRR2153283, SRR2153286, SRR2153287, SRR3102934, SRR3123272, SRR3123273, SRR3123275, SRR3123276, SRR3123277, SRR3123279, SRR3123281, SRR3123337, SRR3123340, SRR3123341, SRR3123342, SRR3123355, SRR3123357, SRR3123359, SRR3123361, SRR3123362, SRR3123364, SRR3123372, SRR3123380, SRR3123385, SRR3123388, SRR3123389, SRR3123390, SRR3123400, SRR3123402, SRR3123404, SRR3123406, SRR3123407, SRR3123408, SRR3123443, SRR3123445, SRR3123446, SRR3123448, SRR3123449, SRR3123451, SRR5041700, SRR5041701, SRR5615334, SRR5615335, SRR5615342, SRR5615343, SRR6047306, SRR6047307, SRR6047308, SRR6047309, SRR6047310, SRR6047311, SRR6047312, SRR6047313, SRR6047314, SRR6047315, SRR6047316, SRR6047317, SRR6727829, SRR6727830, SRR6727831, SRR6727832, SRR6727833, SRR6727834, SRR6727835, SRR6727836, SRR6727837, SRR6727838, SRR6727839, SRR6727840, SRR6727841, SRR6727842, SRR6727843, SRR6727844, SRR6727845, SRR6727846, SRR6727847, SRR6727848, SRR6727849, SRR6727850, SRR6727851, SRR6727852, SRR6727853, SRR6727854, SRR6727855, SRR6727856, SRR6727857, SRR6727858, SRR6727859, SRR6727860, SRR7460786, SRR7460787, SRR7460788, SRR7460789, SRR7460790, SRR7460791, SRR7472351, SRR7472352, SRR7472353, SRR7472354, SRR7472355, SRR7472356, SRR7472357, SRR7472358, SRR7472359, SRR7472360, SRR7472361, SRR7472362 and SRR7472363.

***Blattella germanica*:**
SRR060166, SRR1693414, SRR1693419, SRR1695499, SRR5458588, SRR5458589, SRR5458590, SRR5458591, SRR5458592, SRR5458593, SRR6041830, SRR6041831, SRR9617635, SRR9617636, SRR5656357, SRR5656358, SRR5656359, SRR5656360, SRR5656361, SRR5656362, SRR5656363, SRR5656364, SRR5656365, SRR5656366, SRR5656367, SRR5656368, SRR5656369, SRR5656370, SRR5656371, SRR5656372, SRR6784706, SRR6784707, SRR6784708, SRR6784709, SRR6784710 and SRR6784711.

***Bombus terrestris*:**
ERR883767, ERR883768, ERR883769, ERR883770, ERR883771, ERR883772, ERR883773, ERR883774, ERR883775, ERR883776, ERR883777, ERR883778, ERR883779, ERR883780, ERR883781, ERR883782, ERR883783, ERR883784, ERR883785, ERR883786, ERR883787, ERR883788, ERR883789, ERR883790, ERR883791, ERR883792, ERR883793, SRR5125102, SRR5125103, SRR5125104, SRR5125105, SRR5125106, SRR5125107, SRR5125108, SRR5125109, SRR5125110, SRR5125111, SRR5125112, SRR5125113, SRR5125114, SRR5125115, SRR5125116,

SRR5125117, SRR5125118, SRR5125119, SRR5125120, SRR5125121, SRR5125122, SRR5125123, SRR5125124, SRR5125125, SRR5125126, SRR5125127, SRR5125128, SRR5125129, SRR5125130, SRR5125131, SRR5125132, SRR5125133, SRR5125134, SRR5614374, SRR5614375, SRR5614825, SRR5614826 and SRR5614828.

***Bombyx mori*:**
SRR10035581, SRR10035582, SRR10035583, SRR10035584, SRR10035585, SRR10035586, SRR10035587, SRR10035588, SRR10035589, SRR10035590, SRR10035591, SRR10035592, SRR10035593, SRR10035594, SRR10035595, SRR10035596, SRR10035597, SRR10035598, SRR10035599, SRR10035600, SRR10035601, SRR10035602, SRR10035603, SRR10035604, SRR10035605, SRR10035606, SRR10035607, SRR10035608, SRR10035609, SRR10035610, SRR10035611, SRR10035612, SRR10035613, SRR10035614, SRR10035615, SRR10035616, SRR10035617, SRR10035618, SRR10035619, SRR10035620, SRR10035621, SRR10035622, SRR10035623, SRR10035624, SRR10035625, SRR10035626, SRR10035627, SRR10035628, SRR10035629, SRR10035630, SRR10035631, SRR10035632, SRR10035633, SRR10035634, SRR10035635, SRR10035636, SRR10035637, SRR10035638, SRR10035639, SRR10035640, SRR10035641, SRR10035642, SRR10035643, SRR10035644, SRR10035645, SRR10035646, SRR10035647, SRR10035648, SRR10035649, SRR10035650, SRR10035651, SRR10035652, SRR10035653, SRR10035654, SRR10035655, SRR10035656, SRR10035657, SRR10035658, SRR10035659, SRR10035660, SRR10035661, SRR10035662, SRR10035663, SRR10035664, SRR10035665, SRR10035666, SRR10035667, SRR10035668, SRR10035669, SRR10035670, SRR10035671, SRR10035672, SRR10035673, SRR10035674, SRR10035675, SRR10035676, SRR10035677, SRR10035678, SRR10035679, SRR10035680, SRR10035681, SRR10035682, SRR10035683, SRR10035684, SRR10035685, SRR10035686, SRR10035687, SRR10035688, SRR10035689, SRR10035690, SRR10035691, SRR10035692, SRR10035693, SRR10035694, SRR10035695, SRR10035696, SRR10035697, SRR10035698, SRR10035699, SRR10035700, SRR10035701, SRR10035702, SRR10035703, SRR10035704, SRR10035705, SRR10035706, SRR10035707, SRR10035708, SRR10035709, SRR10035710, SRR10035711, SRR10035712, SRR10035713, SRR10035714, SRR10035715, SRR10035716, SRR10035717, SRR10035718, SRR10035719, SRR10035720, SRR10035721, SRR10035722, SRR10035723, SRR10035724, SRR10035725, SRR10035726, SRR10035727, SRR10035728, SRR10035729, SRR10035730, SRR10035731, SRR10035732, SRR10035733, SRR10035734, SRR10035735, SRR10035736, SRR10035737, SRR10035738, SRR10035739, SRR10035740, SRR10035741, SRR10035742, SRR10035743, SRR10035744, SRR10035745, SRR10035746, SRR10035747, SRR10035748, SRR10035749, SRR10035750, SRR10035751, SRR10035752, SRR10035753, SRR10035754, SRR10035755, SRR10035756, SRR10035757, SRR10035758, SRR10035759, SRR10035760, SRR10035761, SRR10035762, SRR10035763, SRR10035764, SRR10035765, SRR10035766, SRR10035767, SRR10035768, SRR10035769, SRR10035770, SRR10035771, SRR10035772, SRR10035773, SRR10035774, SRR10035775, SRR10035776, SRR10035777, SRR10035778, SRR10035779, SRR10035780, SRR10035781, SRR10035782, SRR10035783, SRR10035784, SRR10035785, SRR10035786, SRR10035787, SRR10035788, SRR10035789, SRR10035790, SRR10035791, SRR10035792, SRR10035793, SRR10035794, SRR10035795, SRR10035796, SRR10035797, SRR10035798, SRR10035799, SRR10035800, SRR10035801, SRR10035802, SRR10035803, SRR10035804, SRR10035805, SRR10035806, SRR10035807, SRR10035808, SRR10035809, SRR10035810, SRR10035811, SRR10035812, SRR10035813, SRR10035814, SRR10035815, SRR10035816, SRR10035817, SRR10035818, SRR10035819, SRR10035820, SRR10035821, SRR10035822, SRR10035823, SRR10035824, SRR10035825, SRR10035826,

SRR10035827, SRR10035828, SRR10035829, SRR10035830, SRR10035831, SRR10035832 and SRR10035833.

***Glossina morsitans***:
SRR1738183, SRR1738184, SRR2424404, SRR2424406, SRR2433816, SRR2433817, SRR2433818, SRR2433820, SRR5207250, SRR5207251, SRR5207252, SRR5838330, SRR5838331, SRR5838332, SRR5838333, SRR5838334, SRR5838335, SRR6943612, SRR6943613, SRR6943614, SRR6943615, SRR8284535, SRR8284536, SRR8284537, SRR8284538, SRR8284539, SRR8284540, SRR8284541, SRR8284543, SRR8284544, SRR8284545, SRR8284546, SRR8284548, SRR8284549, SRR8284550, SRR869502 and SRR869503.

***Gryllus rubens***:
SRR3182687, SRR3182690, SRR3182693, SRR3182696, SRR3182699, SRR3182702, SRR3182705, SRR3182708, SRR3182688, SRR3182691, SRR3182694, SRR3182697, SRR3182700, SRR3182703, SRR3182706, SRR3182709, SRR3182689, SRR3182692, SRR3182695, SRR3182698, SRR3182701, SRR3182704, SRR3182707 and SRR3182710.

***Hermetia illucens***:
ERR1801985, ERR1801986, ERR1801987, ERR1801988, ERR1801989, ERR1801990, ERR1801991, ERR1801992, ERR1801993, ERR1801994, ERR1801995, ERR1801996, ERR1801997, ERR1801998, SRR6656085, SRR6656086, SRR6656087, SRR6656088, SRR10158821 and SRR10233312.

***Latrodectus geometricus***:
SRR5285081, SRR5285082, SRR5285083, SRR5285084, SRR5285085, SRR5285086, SRR5285087, SRR5285088, SRR5285089, SRR5285090, SRR5285091, SRR5285092, SRR5285093, SRR5285094, SRR5285095, SRR5285096, SRR5285097, SRR5285098, SRR5285099 and SRR5285100.

***Latrodectus hesperus***:
SRR1219650, SRR1219651, SRR1219652, SRR1219665, SRR1539570, SRR1853323, SRR1853324, SRR1853325, SRR5285101, SRR5285102, SRR5285103, SRR5285104, SRR5285105, SRR5285106, SRR5285107, SRR5285108, SRR5285109, SRR5285110, SRR5285111, SRR5285112, SRR5285113, SRR5285114, SRR5285115, SRR5285116, SRR5285117, SRR5285118, SRR5285119, SRR5285120, SRR5285121, SRR5285122 and SRR5285123.

***Mesobuthus martensii***:
SRR2592319, SRR2592593, SRR2592957, SRR2592959, SRR2592960, SRR2790072, SRR2791632, SRR2797238, SRR2817451, SRR2829245, SRR3056832, SRR3061371, SRR3061373, SRR3061377, SRR3061379, SRR3984597, SRR3984610, SRR3984642, SRR3984661, SRR3984663 and SRR4188636.

***Parasteatoda tepidariorum***:
SRR1824487, SRR1824488, SRR1824489, SRR5131057, SRR5131058, SRR5602548, SRR5602549, SRR5602550, SRR5602551, SRR6941319, SRR6941320, SRR6941321, SRR6941322, SRR6941323, SRR6941324, SRR6941325, SRR6941326, SRR6941327, SRR6941328, SRR6941329, SRR6941330, SRR6941331, SRR6941332, SRR6941333, SRR6941334, SRR6941335, SRR6941336, SRR6941337, SRR6941338, SRR6941339, SRR6941340, SRR6941341, SRR6941342, SRR6941343, SRR6941344, SRR6941345, SRR6941346, SRR6941347, SRR6941348, SRR6941349, SRR6941350, SRR6941351,

SRR6941352, SRR6941353, SRR6941354, SRR6941355, SRR6941356, SRR6941357, SRR6941358, SRR8755627, SRR8755628, SRR8755629, SRR8755630, SRR8755631, SRR8755632, SRR8755633 and SRR8755634.

***Pediculus humanus*:**
SRR9617639 and SRR9617640.

***Oncopeltus fasciatus*:**
SRR6495591, SRR6495592, SRR6495593, SRR6495594, SRR6495595, SRR6495596, SRR7404757, SRR7404758, SRR7404759, SRR7404760, SRR7404761, SRR7404762, SRR7404763, SRR7404764, SRR7404765, SRR7404766, SRR7404767, SRR7404768, SRR7404769, SRR7404770, SRR7404771, SRR7404772, SRR7404773, SRR7404774, SRR7404775, SRR7404776, SRR7404777, SRR7404778, SRR7404779 and SRR7404780.

***Pardosa pseudoannulata*:**
SRR1970505, SRR1970506, SRR1972491, SRR1972506, SRR1972509, SRR1972511, SRR1972512, SRR1972516, SRR1972520, SRR2010603, SRR2010604, SRR2012296, SRR2012297, SRR2024872, SRR2024873, SRR2024874, SRR2024875, SRR2024876, SRR2024877, SRR6837897, SRR6837898, SRR6837899, SRR6837900, SRR6837901, SRR6837902, SRR6837903, SRR6837904, SRR6837905, SRR7631477, SRR7631478, SRR7631479, SRR7631480, SRR7631481, SRR7631482, SRR7631483, SRR7631484, SRR8083387, SRR8083388, SRR8083389, SRR8083390, SRR8083391, SRR8083392, SRR8083393, SRR8083394, SRR8083395, SRR8083396, SRR8083397 and SRR8083398.

***Periplaneta americana*:**
DRR014884, DRR014885, DRR014886, DRR014887, DRR014888, DRR014889, SRR1184457, SRR1184458, SRR1321695, SRR1322009, SRR2994649, SRR2994650, SRR3056857, SRR3056858, SRR3089536, SRR3089537, SRR3089538, SRR3289663, SRR3289684, SRR3289687, SRR5097509, SRR5097510, SRR5097511, SRR5097512, SRR5097513, SRR5097514, SRR5097515, SRR5097516, SRR5286150, SRR5286151, SRR5286152, SRR5286153 and SRR5286154.

***Rhodnius prolixus*:**
SRR206936, SRR206937, SRR206938, SRR206946, SRR206947, SRR206948, SRR206952, SRR206983, SRR206984, SRR206985, SRR2001240, SRR2001241, SRR2001242, SRR3115162, SRR7738238, SRR7738239, SRR9617637, SRR9617638, ERR1315260, ERR1315262, ERR1315264 and ERR1331764.

***Steatoda grossa*:**
SRR5285124, SRR5285125, SRR5285126, SRR5285127, SRR5285128, SRR5285129, SRR5285130, SRR5285131, SRR5285132, SRR5285133, SRR5285134, SRR5285135, SRR5285136, SRR5285137, SRR5285138, SRR5285139, SRR5285140, SRR5285141 and SRR5285142.

***Stegodyphus dumicola*:**
SRR10216514, SRR10216515, SRR10216516, SRR10216517, SRR10216518, SRR10216519, SRR10216520, SRR10216521, SRR10216522, SRR10216523, SRR10216524 and SRR10216525.

***Tenebrio molitor*:**

DRR002380, SRR1023012, SRR1023013, SRR1023014, SRR1023015, SRR1023016, SRR1023017, SRR1023018, SRR1023019, SRR1023020, SRR1023021, SRR1023022, SRR1023023, SRR1291244 and SRR1636025.

***Timema cristinae*:**
SRR5786829, SRR5786830, SRR5786831, SRR5786832, SRR5786834, SRR5786835, SRR5786915, SRR5786916, SRR5786942, SRR5786943, SRR5786944, SRR5786945, SRR5786946, SRR5786947, SRR5786948, SRR5786949, SRR5786950 and SRR5786951.

***Zootermopsis nevadensis*:**
DRR110536, DRR110537, DRR110538, DRR110539, DRR110540, DRR110541, DRR110542, DRR110543, DRR110544, DRR110545, DRR110546, DRR110547, DRR110548, DRR110549, DRR110550, DRR139981, DRR139982, DRR139983, DRR139984, DRR139985, DRR139986, DRR139987, DRR139988, DRR151559, DRR151560, DRR151561, DRR151562, DRR151563, DRR151564, DRR151565, DRR151566, DRR151567, DRR151568, DRR151569, DRR151570, SRR863596, SRR863597, SRR863598, SRR863599, SRR863601, SRR863602, SRR863603, SRR863604, SRR863605, SRR863606, SRR863612, SRR863613, SRR863614, SRR1167035, SRR1167037, SRR1167039, SRR1167040, SRR1167041, SRR1167042, SRR1167043, SRR1167044, SRR1167178, SRR1167247, SRR1167255, SRR1167256, SRR3139733, SRR3139734, SRR3139735, SRR3139736, SRR3139737, SRR3139738, SRR3139739, SRR3139740, SRR3139741, SRR3139742, SRR3139743 and SRR4240472.

On the next three pages:

**Fig. S1.** Sequence similarity tree of arthropod insulin-like peptides. Ilp sequences identified in this manuscript and a previous one (Veenstra, 2020). It is a different representation of figure 3 of the main manuscript with identification of the different sequences. Branch probabilities have only been indicated for the major clusters.

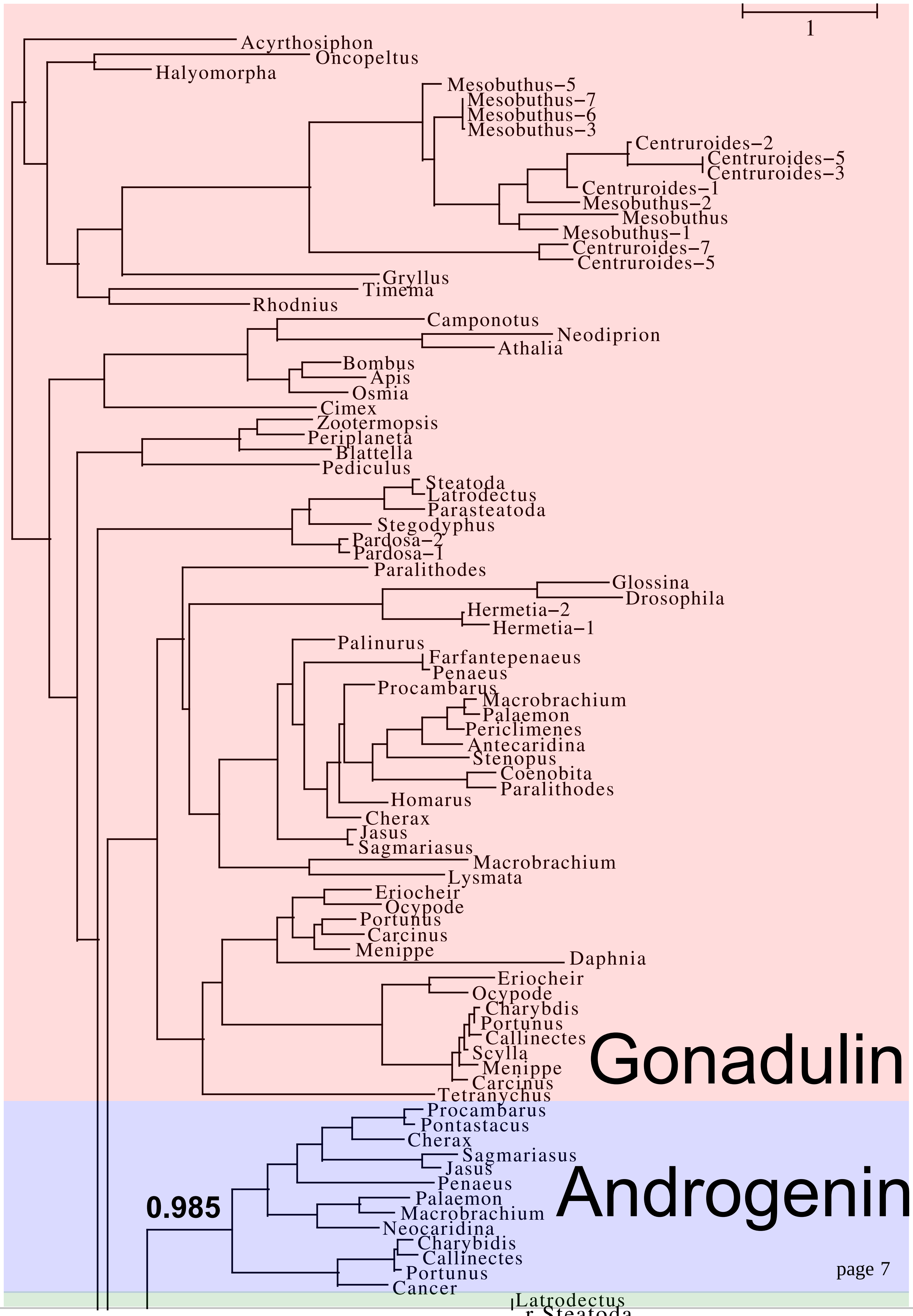
1
Acyrthosiphon
Oncopeltus
Halyomorpha
Mesobuthus−5
Mesobuthus−7
Mesobuthus−6
Mesobuthus−3
Centruroides−2
Centruroides−5
Centruroides−3
Centruroides−1
Mesobuthus−2
Mesobuthus
Mesobuthus−1
Centruroides−7
Centruroides−5
Gryllus
Timema
Rhodnius
Camponotus
Neodiprion
Athalia
Bombus
Apis
Osmia
Cimex
Zootermopsis
Periplaneta
Blattella
Pediculus
Steatoda
Latrodectus
Parasteatoda
Stegodyphus
Pardosa−2
Pardosa−1
Paralithodes
Glossina
Drosophila
Hermetia−2
Hermetia−1
Palinurus
Farfantepenaeus
Penaeus
Procambarus
Macrobrachium
Palaemon
Periclimenes
Antecaridina
Stenopus
Coenobita
Paralithodes
Homarus
Cherax
Jasus
Sagmariasus
Macrobrachium
Lysmata
Eriocheir
Ocypode
Portunus
Carcinus
Menippe
Daphnia
Eriocheir
Ocypode
Charybdis
Portunus
Callinectes
Scylla
Menippe
Carcinus
Tetranychus
Gonadulin
Procambarus
Pontastacus
Cherax
Sagmariasus
Jasus
Penaeus
Palaemon
Macrobrachium
Neocaridina
Charybidis
Callinectes
Portunus
Cancer
0.985
Androgenin
Latrodectus

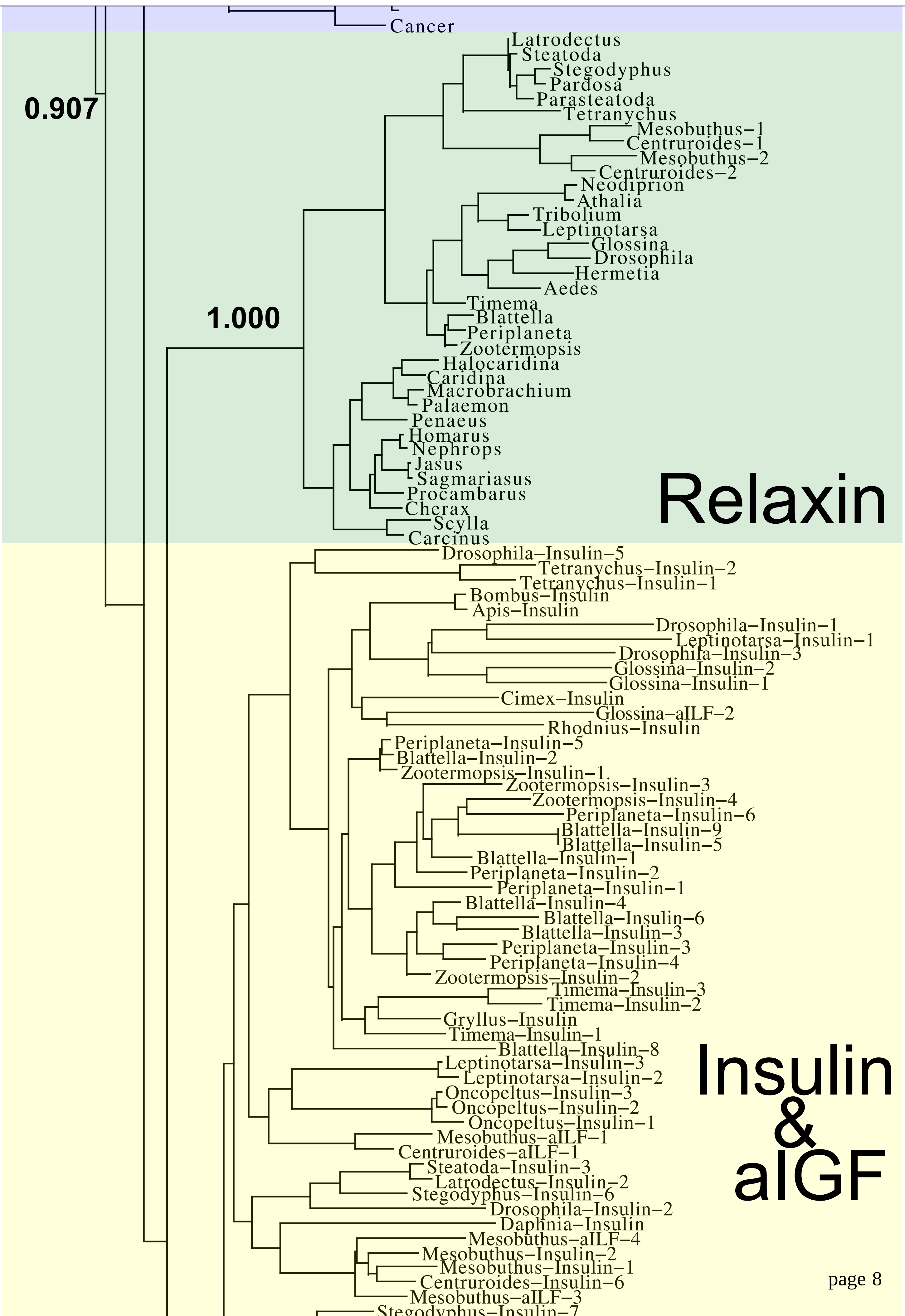
Cancer
0.907
Latrodectus
Steatoda
Stegodyphus
Pardosa
Parasteatoda
Tetranychus
Mesobuthus−1
Centruroides−1
Mesobuthus−2
Centruroides−2
Neodiprion
Athalia
Tribolium
Leptinotarsa
Glossina
Drosophila
Hermetia
Aedes
Timema
1.000
Blattella
Periplaneta
Zootermopsis
Halocaridina
Caridina
Macrobrachium
Palaemon
Penaeus
Homarus
Nephrops
Jasus
Sagmariasus
Procambarus
Cherax
Scylla
Carcinus
Relaxin
Drosophila−Insulin−5
Tetranychus−Insulin−2
Tetranychus−Insulin−1
Bombus−Insulin
Apis−Insulin
Drosophila−Insulin−1
Leptinotarsa−Insulin−1
Drosophila−Insulin−3
Glossina−Insulin−2
Glossina−Insulin−1
Cimex−Insulin
Glossina−aILF−2
Rhodnius−Insulin
Periplaneta−Insulin−5
Blattella−Insulin−2
Zootermopsis−Insulin−1
Zootermopsis−Insulin−3
Zootermopsis−Insulin−4
Periplaneta−Insulin−6
Blattella−Insulin−9
Blattella−Insulin−5
Blattella−Insulin−1
Periplaneta−Insulin−2
Periplaneta−Insulin−1
Blattella−Insulin−4
Blattella−Insulin−6
Blattella−Insulin−3
Periplaneta−Insulin−3
Periplaneta−Insulin−4
Zootermopsis−Insulin−2
Timema−Insulin−3
Timema−Insulin−2
Gryllus−Insulin
Timema−Insulin−1
Blattella−Insulin−8
Leptinotarsa−Insulin−3
Leptinotarsa−Insulin−2
Oncopeltus−Insulin−3
Oncopeltus−Insulin−2
Oncopeltus−Insulin−1
Mesobuthus−aILF−1
Centruroides−aILF−1
Steatoda−Insulin−3
Latrodectus−Insulin−2
Stegodyphus−Insulin−6
Drosophila−Insulin−2
Daphnia−Insulin
Mesobuthus−aILF−4
Mesobuthus−Insulin−2
Mesobuthus−Insulin−1
Centruroides−Insulin−6
Mesobuthus−aILF−3
Stegodyphus−Insulin−7
Insulin
&
aIGF

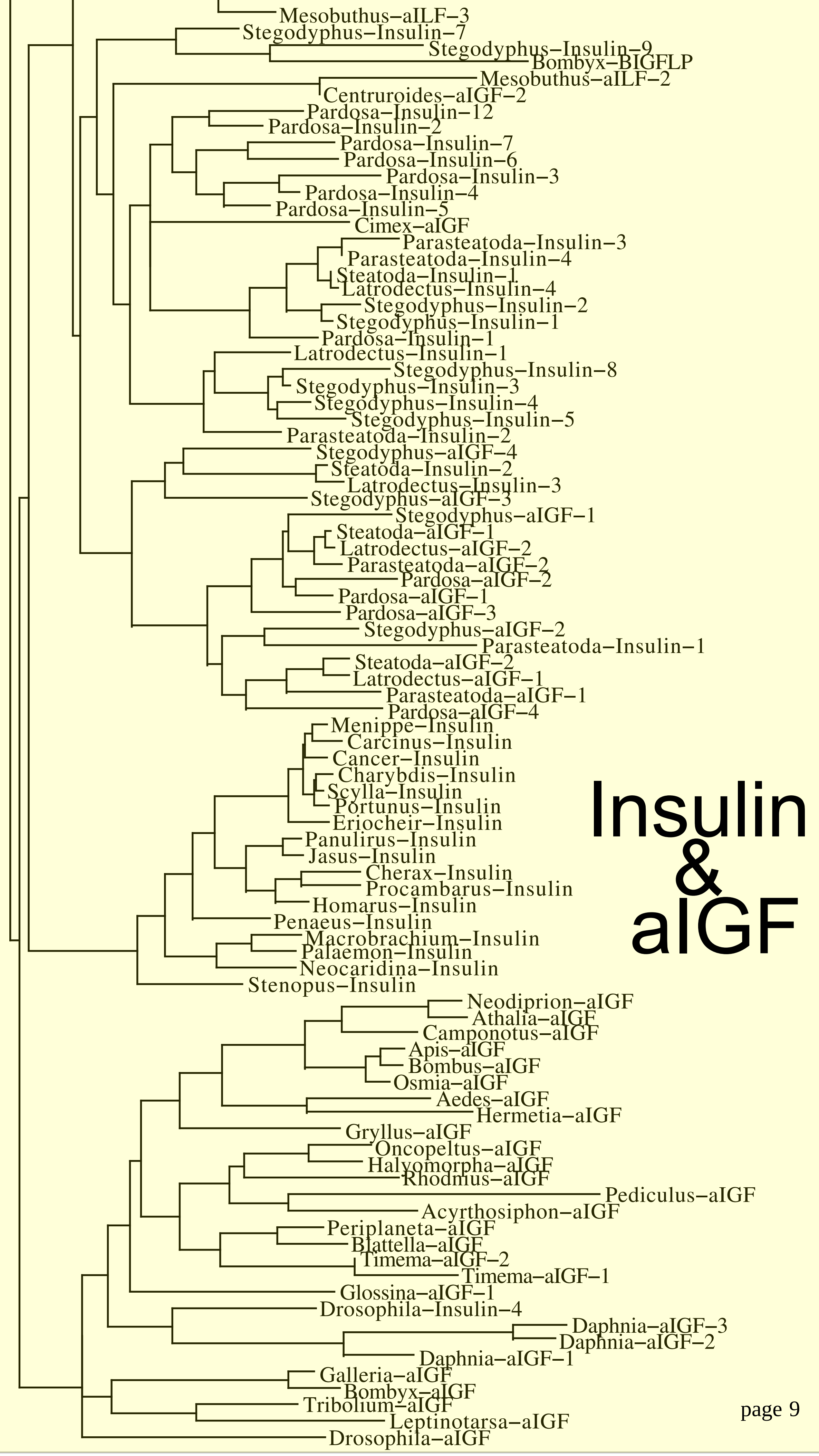
Mesobuthus–aILF–3
Stegodyphus–Insulin–7
Stegodyphus–Insulin–9
Bombyx–BIGFLP
Mesobuthus–aILF–2
Centruroides–aIGF–2
Pardosa–Insulin–12
Pardosa–Insulin–2
Pardosa–Insulin–7
Pardosa–Insulin–6
Pardosa–Insulin–3
Pardosa–Insulin–4
Pardosa–Insulin–5
Cimex–aIGF
Parasteatoda–Insulin–3
Parasteatoda–Insulin–4
Steatoda–Insulin–1
Latrodectus–Insulin–4
Stegodyphus–Insulin–2
Stegodyphus–Insulin–1
Pardosa–Insulin–1
Latrodectus–Insulin–1
Stegodyphus–Insulin–8
Stegodyphus–Insulin–3
Stegodyphus–Insulin–4
Stegodyphus–Insulin–5
Parasteatoda–Insulin–2
Stegodyphus–aIGF–4
Steatoda–Insulin–2
Latrodectus–Insulin–3
Stegodyphus–aIGF–3
Stegodyphus–aIGF–1
Steatoda–aIGF–1
Latrodectus–aIGF–2
Parasteatoda–aIGF–2
Pardosa–aIGF–2
Pardosa–aIGF–1
Pardosa–aIGF–3
Stegodyphus–aIGF–2
Parasteatoda–Insulin–1
Steatoda–aIGF–2
Latrodectus–aIGF–1
Parasteatoda–aIGF–1
Pardosa–aIGF–4
Menippe–Insulin
Carcinus–Insulin
Cancer–Insulin
Charybdis–Insulin
Scylla–Insulin
Portunus–Insulin
Eriocheir–Insulin
Panulirus–Insulin
Jasus–Insulin
Cherax–Insulin
Procambarus–Insulin
Homarus–Insulin
Penaeus–Insulin
Macrobrachium–Insulin
Palaemon–Insulin
Neocaridina–Insulin
Stenopus–Insulin
Neodiprion–aIGF
Athalia–aIGF
Camponotus–aIGF
Apis–aIGF
Bombus–aIGF
Osmia–aIGF
Aedes–aIGF
Hermetia–aIGF
Gryllus–aIGF
Oncopeltus–aIGF
Halyomorpha–aIGF
Rhodnius–aIGF
Pediculus–aIGF
Acyrthosiphon–aIGF
Periplaneta–aIGF
Blattella–aIGF
Timema–aIGF–2
Timema–aIGF–1
Glossina–aIGF–1
Drosophila–Insulin–4
Daphnia–aIGF–3
Daphnia–aIGF–2
Daphnia–aIGF–1
Galleria–aIGF
Bombyx–aIGF
Tribolium–aIGF
Leptinotarsa–aIGF
Drosophila–aIGF
Insulin & aIGF

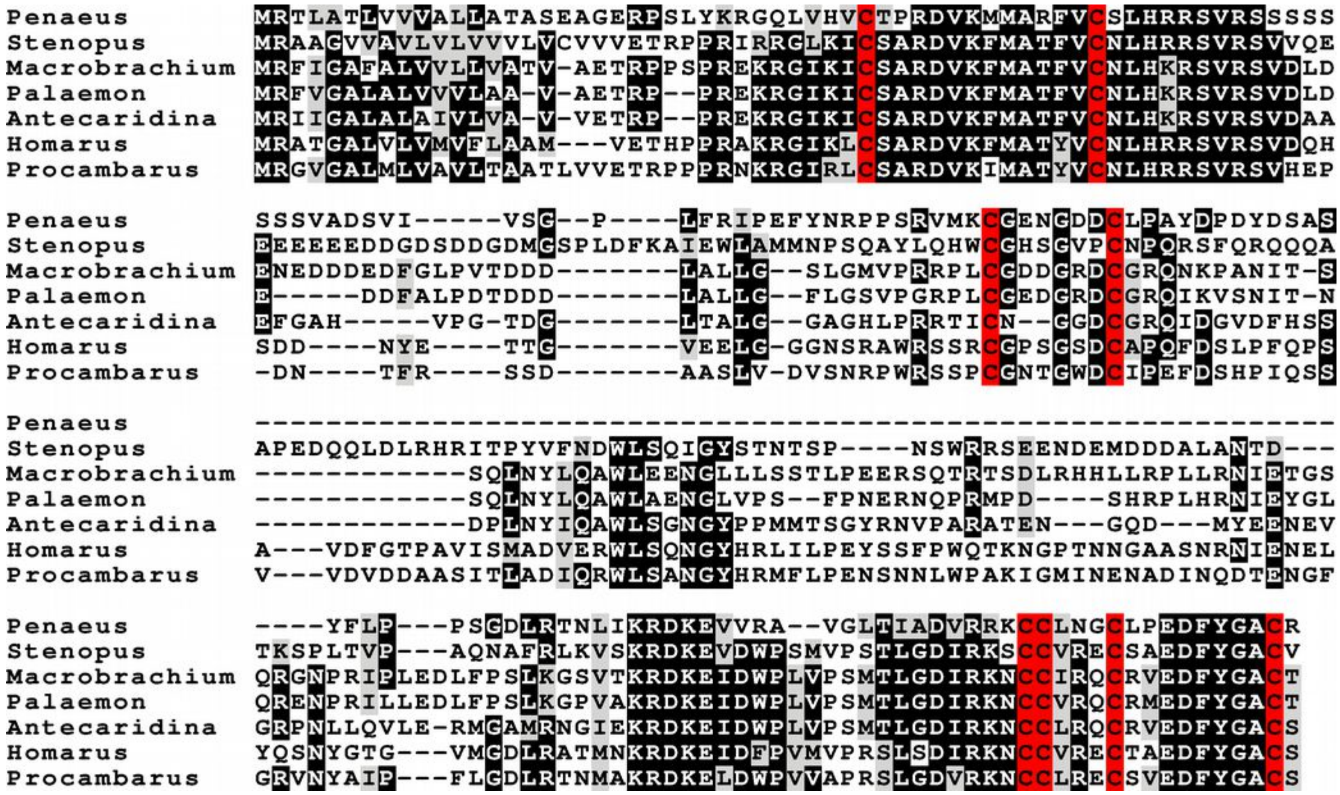


**Fig. S2.** Sequence alignment of decapod gonadulin 2 precursors that have four predicted disulfide bridges. Sequences from Veenstra, 2020. Cysteine residues are in red, conserved amino acid residues in black and conservative substitutions are in grey.

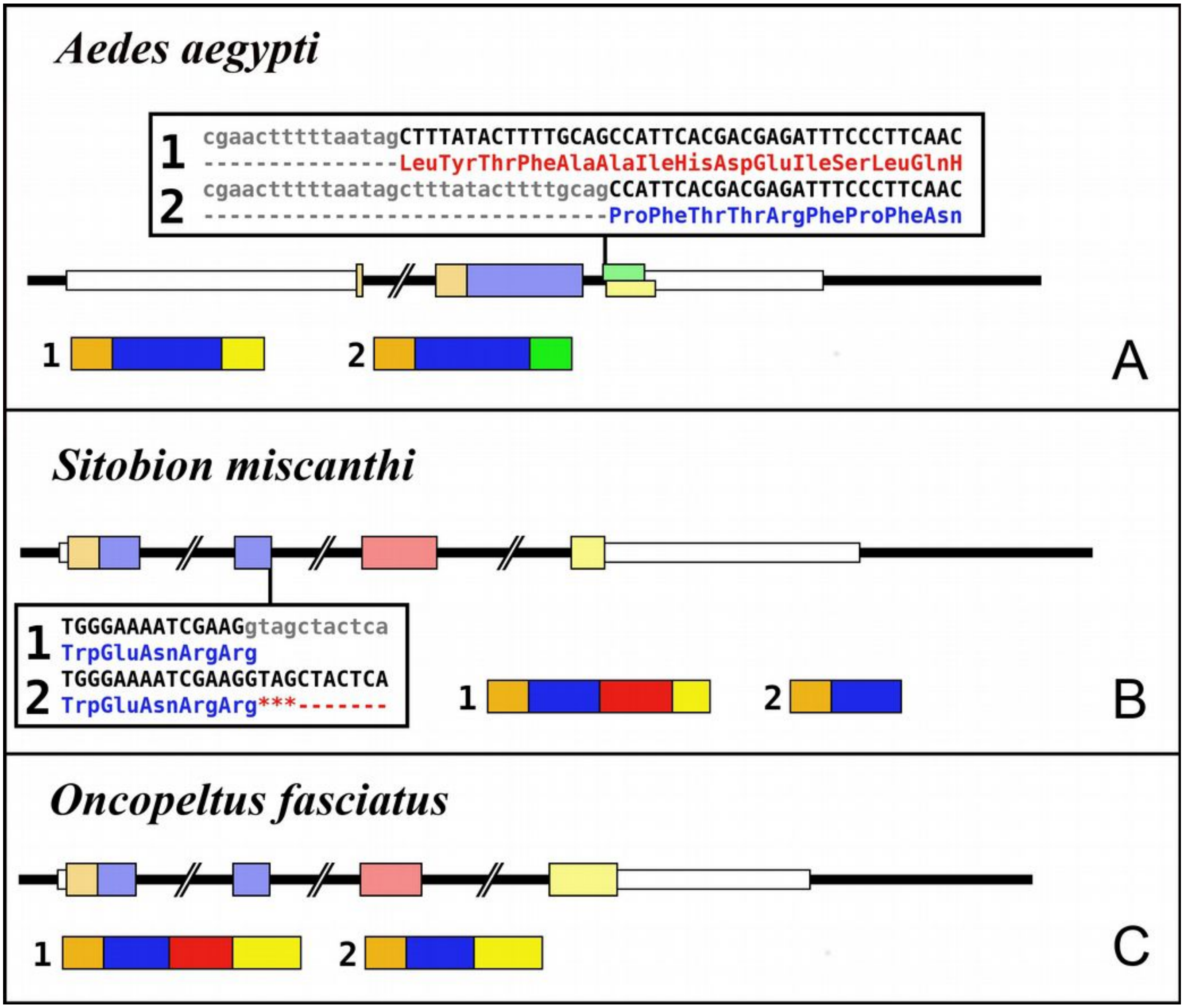


**Fig. S3.** Insect aIGF gene structures. Intron-exon structures of the aIGF genes from *Aedes aegypti* (A), *Sitobion miscanthi* (B) and *Oncopeltus fasciatus* (C). In the top part for each species the gene structure with the exon represented as small colored boxes and the introns as a black line. The orange brown color in the first exon corresponds to the coding sequence of the signal peptide and the blue corresponds to the insulin sequence, while the red color corresponds to the coding sequence of the arginine-rich region that is alternatively spliced in the two isoforms produced from these genes. The yellow exons contains coding sequence for GTVX$_1$PX$_2$(F/Y) consensus sequence. The amino acid sequence coded by this alternatively spliced DNA sequence is indicated. The numbers 1 and 2 show the coding sequences of the two mRNA species produced from these gene using the same colors as for the gene structures. Note that the *Sitobion* and *Oncopeltus* genes are quite similar to the archetype insect aIGF gene, except that the alternative splicing concerns the complete third coding exon and not just the second half of it. The *Aedes* gene on the other hand has lost the third coding exon, but acquired a novel alternative splicing site in what used to be the fourth coding exon. One these splice forms codes for the GTVX$_1$PX$_2$(F/Y) consensus sequence. The *Sitobion* gene was deduced using a transcript from *S. avenae* (GAPL01021540.1).

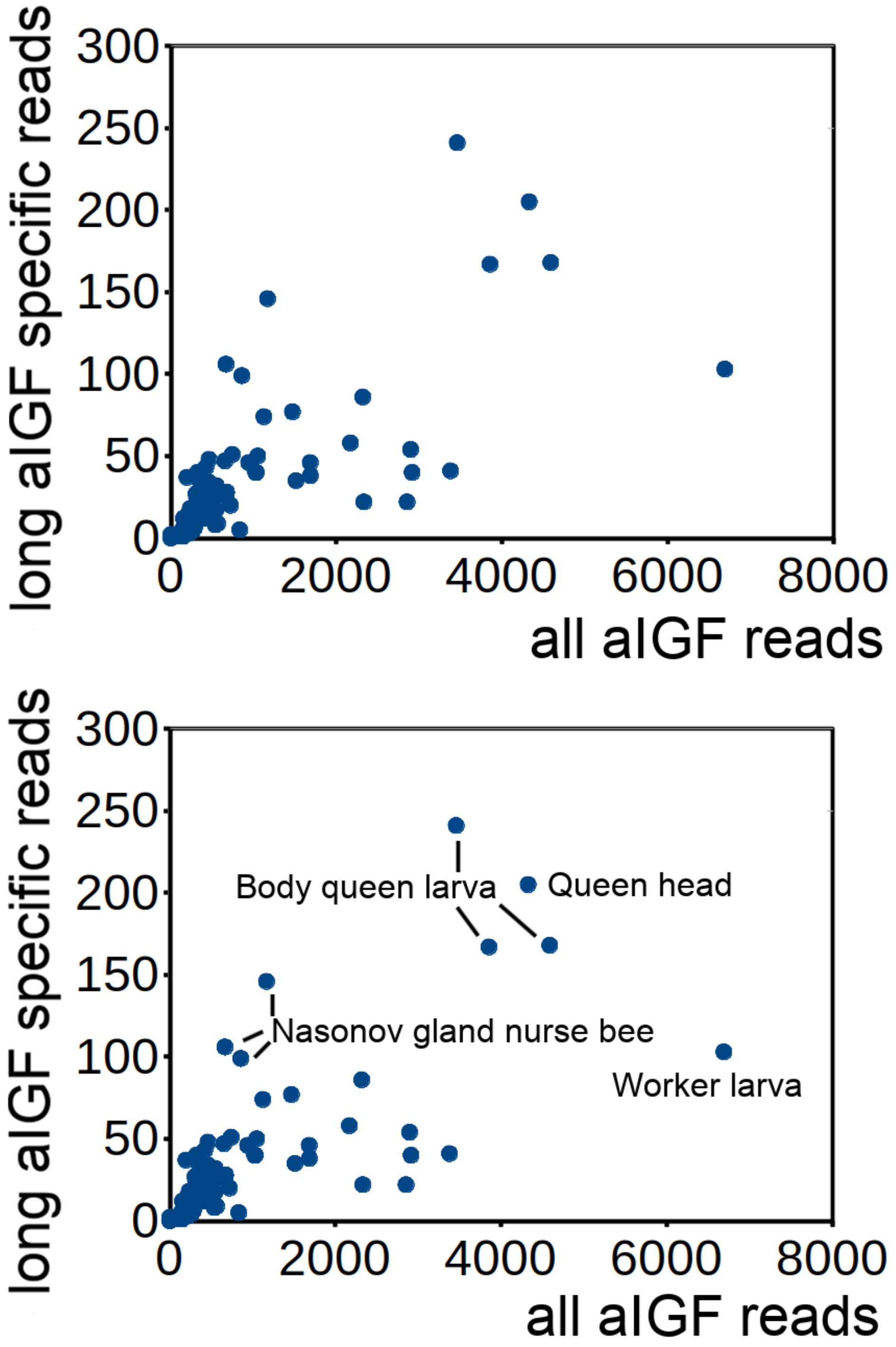


**Fig. S4.** Altenative splicing of *Apis mellifera* aIGF. Correlation between the number of RNAseq reads specific for the long isoform of aIGF and all aIGF reads in honeybee transcriptome SRAs. Top shows the raw results illustrating that the ratio is quite variable. Bottom shows the same data with the source of mRNA that show relatively high expression of the long isoform. The queen larve body data are from 4-day old animals. Other samples can be identified from Suppl. Spreadsheet 2.

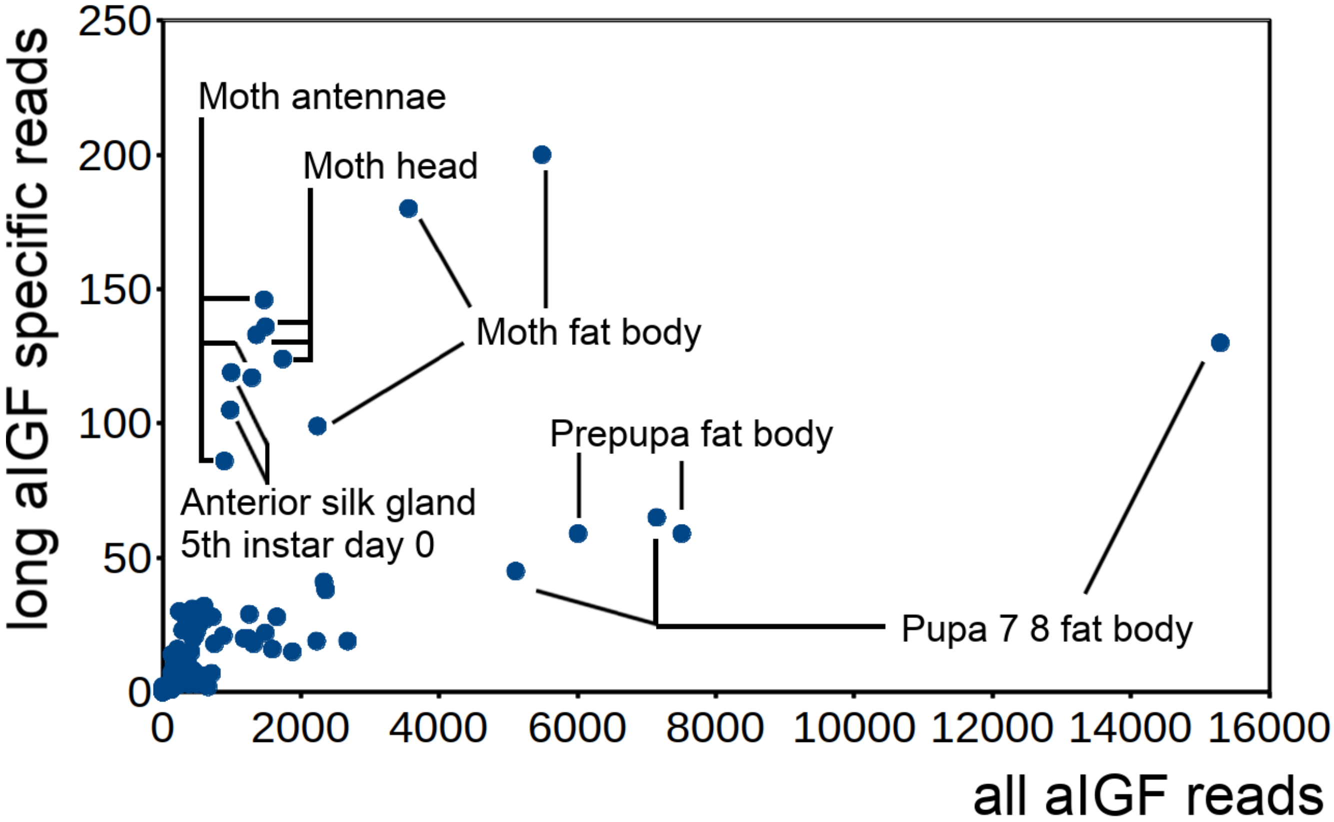


**Fig. S5.** Same data as in figure 7, but with identification of samples that show relatively large numbers for all aIGF transcripts and/or long aIGF transcript specific reads. Note that samples obtained from 1-day old moths contain large numbers of long isoform specific reads, as do anterior silk gland samples form day 0 of the $5^{th}$ instar. Other samples can be identified from Suppl. Spreadsheet 2.

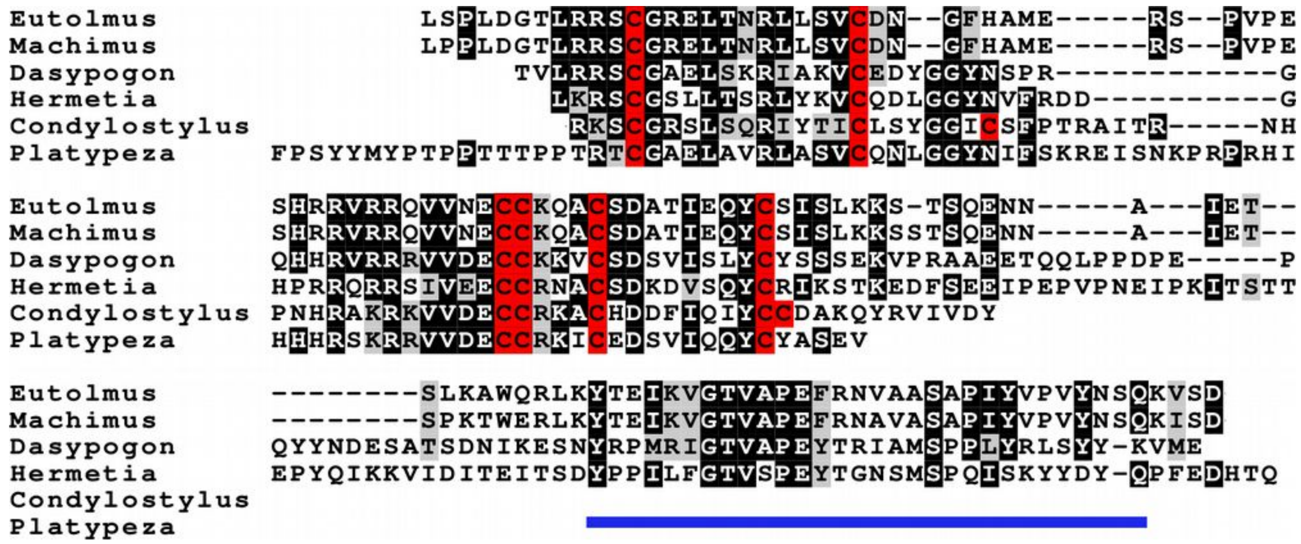


**Fig. S6.** Sequence alignment of aIGFs from three asilids (robber flies) and one stratiomyids (soldier fly), the most evolved non-Eremoneura flies together with one dolichopodid (long-legged fly) and a platypezid (flat-footed fly). The latter two are the least evolved Eremoneura. It is clear that the robberflies (*Eutolmus, Machimus* and *Dasypogon*) and the soldier fly (*Hermetia*) still have the exon coding for the GTV$X_1$P$X_2$(F/Y) consensus sequence (underlined in blue), while both the long-legged fly (*Condylostylus*) and the flat-footed fly (*Platypeza*) have lost this sequences. The remainder of these sequences shows these genes to orthologs. Interestingly, the *Condylostylus* sequence has an additional disulfide bridge. Cysteine residues are in red, conserved amino acid residues in black and conservative substitutions are in grey. Sequences used: *Eutolmus rufibarbis* (GFGA01018701.1); *Dasypogon diadema* (GFGB01021966.1); *Machimus arthriticus* (GFZQ01007432.1); *Hermetia illucens* (Suppl. Spreadsheet 1); *Platypeza anthrax* (GCGU01007956.1). The incomplete sequence of *Condylostylus patibulatus* was deduced from transcriptome SRAs SRR5559329 and SRR5559330 and confirmed by the genomic reads from SRAs SAMN03220540 and SAMN03220541. Importantly, the extra cysteine residues were found in both the transcriptome and genom reads, while the stop codon in the transcriptome reads was confirmed by in the genome sequences that do not show a intron splice site.

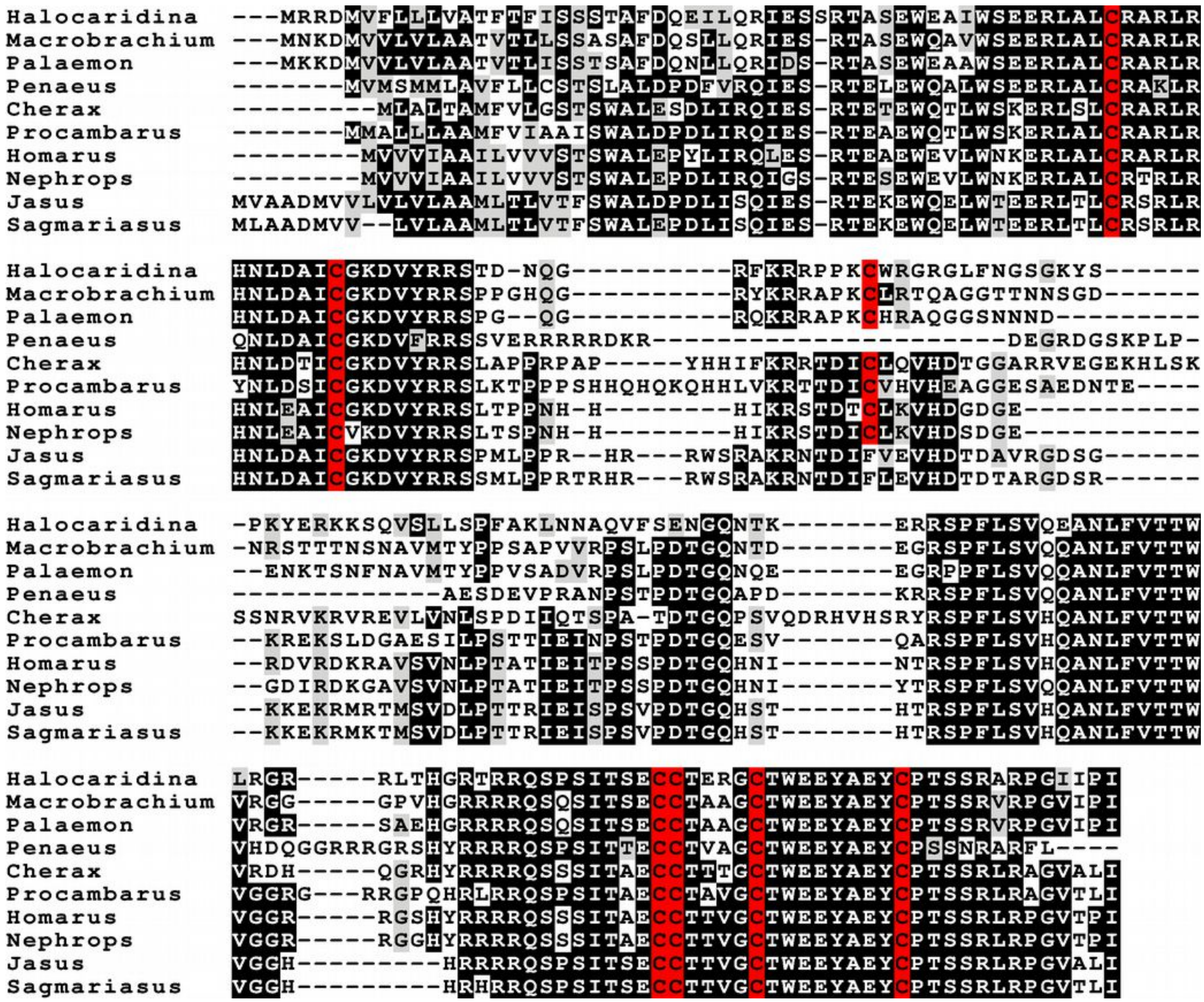

**Fig. S7.** Sequence alignment of decapod relaxin precursors. Several of the predicted proteins have seven cysteine residues, which is unexpected as the cysteine residues form disulfide bridges, but in order to do so one needs an even number. Cysteine residues are in red, conserved amino acid residues in black and conservative substitutions are in grey.
